# Validation of an optimised Oxford Nanopore sequencing workflow versus Illumina for mycobacteria from primary MGIT culture

**DOI:** 10.64898/2026.02.04.703726

**Authors:** Catriona S Baker, Matthew Colpus, Jess Gentry, Alexandra Hall, Eloïse Roghi, Hermione Webster, Bronte Drummond, Ruth Cooper, Hieu Thai, Jeremy Westhead, Robert Turner, Timothy EA Peto, Philip W Fowler, Marcus Morgan, Derrick W Crook

**Affiliations:** Modernising Medical Microbiology Unit, Nuffield Department of Medicine, University of Oxford; Oxford University Hospitals NHS Foundation Trust; National Institute of Health Research Oxford Biomedical Research Centre, University of Oxford, Oxford, UK; Health Protection Research Unit in Healthcare Associated Infections and Antimicrobial Resistance, John Radcliffe Hospital, Oxford, U.K; Shared Hospital Laboratory and Sunnybrook Research Institute, Sunnybrook Hospital, Toronto, Canada

**Keywords:** Mycobacteria, whole genome sequencing, DNA extraction, long read sequencing

## Abstract

Illumina sequencing of primary MGIT™ cultures is an established workflow in several reference mycobacteriology laboratories. Oxford Nanopore Technologies (ONT) provides real-time genetic sequencing yielding long reads which help resolve repetitive genomes and is being explored for in-house implementation within diagnostic laboratories. However, low DNA yields from primary MGIT cultures frequently limit the application of ONT workflows, due to high minimum DNA input requirements for library preparation.

We validated a modified ONT workflow combining rapid, semi-automated DNA extraction from MGIT cultures with Rapid PCR-Barcoding for whole-genome amplification, and compared its performance with Illumina sequencing for species identification and *Mycobacterium tuberculosis* complex (MTBC) single-nucleotide polymorphism (SNP) detection. A platform-agnostic analysis pipeline enabled consistent human read removal, taxonomic assignment, and MTBC genomic characterisation. ONT sequencing data was subsampled at 1, 6, and 72 hours to determine the earliest time point for reliable species identification.

The concordance between sequencing platforms on species classification was 98.3% (95.8-99.5%) with all differences arising from potential mixed infections. SNP agreement was high, with a mean of 0.3 and a median of 0 SNP differences between sequencing platforms after masking. These findings demonstrate the feasibility of PCR-amplified ONT sequencing as a reliable alternative for routine genomic characterisation of MGIT cultures.

**IMPORTANCE:** Rapid identification of mycobacterial infections is essential for timely patient care and infection control. Many clinical laboratories currently rely on outsourcing sequencing to external reference centres, which adds time and delays the return of clinically actionable results. These services commonly use Illumina sequencing, which, while highly accurate, involves complex workflows and longer turnaround times. Newer technologies offer the potential to generate results in real time, but their use has been limited by the low amount of DNA available from routine culture samples.

In this study, we developed an improved workflow that increases the amount of usable DNA and enables reliable, rapid sequencing directly from these samples. This approach allows multiple samples to be processed together and reduces the time needed to obtain results. Importantly, it could enable clinical laboratories to perform sequencing in-house, reducing reliance on external services and improving turnaround times.

## INTRODUCTION

The genus *Mycobacterium* comprises more than 190 recognized species, many of which are capable of causing a wide range of human diseases, including pulmonary, cutaneous, and systemic infections^1,2^. Globally, the *Mycobacterium tuberculosis* complex (MTBC) remains a leading cause of infectious disease, with an estimated 10.7 million new cases and 1.23 million deaths worldwide due to tuberculosis (TB) in 2024^3^. In addition, non-tuberculous mycobacteria (NTM) are emerging in prominence as opportunistic pathogens, particularly among older individuals and those with impaired host defences, with associated mortality more than doubling in the last two decades^4^.

Historically, *Mycobacterium* species were identified by phenotypic and biochemical assays, later supplemented by commercial nucleic acid probe-based assays, such as the Hain Lifescience GenoType Mycobacterium CM assay^5,6^. As many of these organisms are characteristically slow-growing, traditional diagnostic methods often require several days to weeks to achieve species-level identification and could still yield inconclusive results, particularly when differentiating closely related species or detecting mixed infections ^7^. More recently, species identification has improved considerably with the adoption of modern platforms such as MALDI-TOF mass spectrometry, which is able to rapidly and accurately identify many NTM species ^8^. However, for *Mycobacterium spp*. and the MTBC in particular, whole-genome sequencing (WGS) offers greater resolution from a single assay, enabling precise species differentiation, lineage assignment, and antimicrobial-resistance (AMR) prediction beyond the capabilities of MALDI-TOF. In addition, WGS resolves single-nucleotide polymorphisms (SNPs), facilitating faster and more accurate detection of transmission clusters and outbreaks ^9^.

The implementation of WGS in clinical settings is often constrained by upstream challenges in sample preparation. DNA extraction from mycobacterial cultures using conventional methods such as cetyltrimethylammonium bromide (CTAB) is lengthy, labour-intensive, and poorly suited to high-throughput or automated workflows. To address these limitations, we developed and optimised a more scalable and automatable cell lysis and DNA extraction method.

Following extraction, DNA can be sequenced using short-read platforms such as Illumina or MGI Tech, or long-read platforms such as Oxford Nanopore Technologies (ONT) or Pacific Biosciences (PacBio). Illumina sequencing remains the most widely adopted approach due to its maturity and high base-calling accuracy and is often regarded as the gold standard when assessing newer technologies ^10^. This accuracy achieved by Illumina platforms confers a major advantage for reliable SNP detection^11^. However, Illumina sequencing typically generates reads of up to 300 bp, which complicates the resolution of repetitive or complex genomic regions (e.g. the PE/PPE gene family in *M. tuberculosis*), potentially leading to incomplete genomic assemblies ^12,13^.

Long-read sequencing with Oxford Nanopore Technologies (ONT) offers several technical advantages, including real-time data generation and the potential for live analysis, allowing sequencing runs to be monitored and terminated once sufficient data have been obtained. In addition, the generation of long reads can improve the resolution of repetitive genomic regions compared with conventional short-read sequencing approaches.

Nonetheless, sequencing mycobacterial DNA from primary MGIT cultures using ONT remains challenging, as DNA yields are frequently insufficient for current ONT library-preparation kits, which typically recommend input quantities exceeding 200 ng per sample. For ONT sequencing, insufficient input DNA frequently results in low coverage and depth, limiting species identification and preventing reliable AMR prediction^10,14,15^. These constraints necessitate the use of PCR-based ONT library preparation strategies when working with low-yield MGIT-derived DNA.

Recent improvements in ONT chemistry and hardware, particularly the R10.4.1 flow cells with V14 chemistry, have improved base-calling accuracy, bringing ONT performance closer to that of Illumina ^16,17^.

A streamlined ONT-compatible workflow for sequencing mycobacterial isolates directly from MGIT cultures has not yet been described, limiting the adoption of ONT for routine clinical mycobacterial genomics and underscoring the need for efficient DNA extraction and amplification methods tailored to ONT sequencing. In this study, we describe the development and validation of a semi-automated DNA extraction workflow and a modified rapid PCR barcoding library preparation protocol optimised for use with low-yield MGIT-derived DNA for ONT sequencing. We assess whether this ONT-based workflow can deliver accurate *Mycobacterium* species identification, SNP detection, and AMR gene profiling for *M. tuberculosis*, with performance comparable to Illumina sequencing, and evaluate the sequencing time required to achieve adequate diagnostic yield.

## MATERIALS AND METHODS

### Sample processing

Samples submitted to the Oxford University Hospitals (OUH) microbiology laboratory for detection of *Mycobacterium* spp. were processed according to specimen type. Non-sterile specimens underwent decontamination using sodium hydroxide (NaOH), whereas sterile specimens were cultured without prior decontamination. Following processing, specimens were inoculated into MGIT™ tubes (Becton Dickinson) supplemented with PANTA (Polymyxin B, Amphotericin B, Nalidixic Acid, Trimethoprim, and Azlocillin) and incubated at 37°C for up to 56 days. Detection of mycobacterial growth was monitored using the MGIT system, and positivity was confirmed by Ziehl–Neelsen staining to demonstrate the presence of acid-fast bacilli (AFB).

An aliquot of 1 mL from each positive MGIT tube was stored at 2 - 8 °C for up to 5 days prior to batching for further processing. Samples were then incubated at 37 °C for 5 days to enhance growth, centrifuged at 6,000g for 5 minutes, and the resulting pellet was resuspended in 200 µL phosphate-buffered saline (PBS; EO Labs) in a 1.5 mL Sarstedt screw-top tube.

Each sample was analysed separately, and multiple samples from the same patient were included where available (i.e. patient samples were not de-duplicated). Duplicate samples collected within different time frames were available for three patients.

### Heat inactivation and validation

Samples were heat inactivated in a fan-assisted oven at 80 ° C for 30 minutes. After heat inactivation, samples were transferred to a Containment Level 2 laboratory. Heat inactivation of *Mycobacterium spp*. was validated by OUH Microbiology, using 20 *Mycobacterium* isolates in MGIT fluid processed as described above, with no growth detected following 56 days of incubation.

### Cell lysis

0.08-0.1g Lysing Matrix B beads (MP Biomedicals, CA, USA) were added to 200 µL heat inactivated sample before undergoing two rounds of mechanical lysis using the MP Bio FastPrep 24 (MP Biomedicals, CA, USA) at 6.0m/s for 40s with a 5-minute interval between lysis rounds. Lysed supernatant was collected following centrifugation at 16,000g for 10 mins at room temperature.

### DNA purification

Lysed supernatant underwent DNA purification using the KingFisher™ Flex (Thermo Scientific ™) system with the addition of 1X AMPure XP beads (Beckman Coulter, CA, USA) and washed twice with 70% ethanol and then eluted into 50 µL elution buffer (CDT-01, Quickgene).

### Quantification

Extracted DNA was measured using a Qubit™ 4.0 fluorometer with the 1X High Sensitivity dsDNA kit, or for samples *>* 120 ng*/*µL, the 1X Broad Range dsDNA kit (Invitrogen, CA, USA).

Illumina sequencing was commissioned from a commercial supplier, GENEWIZ Germany GmbH (Leipzig, Germany), which recommends a minimum DNA concentration of 6 ng*/*µL for its Whole Genome Sequencing (human/non-human) library preparation workflow. For ONT sequencing, the Rapid PCR Barcoding workflow requires a minimum DNA input of 1 ng in 3 µL (equivalent to 0.334 ng*/*µL). To enable paired ONT–Illumina comparative analyses while maximising sample inclusion, an internal DNA concentration threshold of 1 ng*/*µL was applied, and samples below this threshold were excluded from downstream analyses.

### Sequencing

#### Illumina

Short-read sequencing was performed by a commercial provider as described above. Library preparation was performed using the NEBNext® Ultra™ II DNA Library Preparation Kit, and Illumina clustering and sequencing were carried out in accordance with the manufacturer’s instructions.

Briefly, genomic DNA was fragmented by acoustic shearing using a Covaris instrument, followed by end repair and 3^*′*^-end adenylation. Adapters were ligated, followed by enrichment by limited-cycle PCR. Libraries were validated using the Agilent 5600 Fragment Analyzer and quantified using a Qubit 4.0 Fluorometer.

Validated libraries were multiplexed and loaded onto an Illumina NovaSeq X Plus instrument according to the manufacturer’s instructions. Sequencing was performed using a 2 × 150 bp paired-end configuration. Image analysis and base calling were performed using NovaSeq Control Software. Raw sequencing data (.bcl files) were converted to FASTQ files and demultiplexed using Illumina bcl2fastq software.

#### ONT

ONT sequencing was performed using a PCR-based rapid barcoding workflow rather than a PCR-free protocol, in accordance with manufacturer recommendations for low-input DNA samples. Long-read sequencing was performed using a modified version of the ONT Rapid PCR Barcoding Kit (SQK-RPB114.24) on R10.4.1 flow cells using v14 chemistry.

The protocol was modified by substituting in Ex Premier™ DNA Polymerase (TaKaRa) for the recommended LongAmp Hot Start Taq 2× Master Mix, as the latter yielded inconsistent PCR amplification. PCR amplification was performed using the cycling conditions detailed in Table 1, with 30 cycles of denaturation, annealing, and extension.

**Table 1:** PCR cycling conditions. 30 cycles of denaturation, annealing, and extension were performed.

| Step | Temperature (°C) | Time (Min:Sec) |  |
| --- | --- | --- | --- |
| Initial | 94 | 1:00 |  |
| Denature | 98 | 0:10 | 30 cycles |
| Anneal | 56 | 0:15 |  |
| Extend | 68 | 6:00 |  |
| Final extension | 65 | 6:00 |  |
|  | 4 | Hold |  |

Except for this modification, standard procedures for the SQK-RPB114.24 kit were followed. SUP basecalling was used with Dorado 4.3.0 using the onboard MinKNOW on an ONT GridION.

#### Controls

For both ONT and Illumina sequencing, *Mycobacterium tuberculosis* H37Rv was included as a control (Illumina n=4, ONT n=2). One *Mycobacterium bovis* Bacillus Calmette-Guérin (BCG) Pasteur strain sample was additionally sequenced using ONT. H37Rv and BCG were obtained from reference stocks held at MMM.

#### Bioinformatics

Sequencing data was processed using an instance of an online cloud platform ^18^. In brief, all samples had human reads removed using Hostile (v1.1.0) or Deacon (v0.7.0) ^19,20^, before quality filtering with Fastp (v0.24.1) ^21^. Kraken2 (v2.1.3) was used to select reads from the *Mycobacteriaceae* family (using standard index 6/5/2023) ^22^. Sylph (v0.9.0) was used after Kraken2 for species identification using the GTDB taxonomy^23^. Reads were then mapped using Minimap2 (v2.28 with ‘map-ont’ preset) against a multifasta of *Mycobacterium* reference genomes to select for MTBC reads and to provide an alternative method of mycobacterial species identification (competitive mapping), whilst Mykrobe (v0.13.0) was used to determine lineage^24,25^.

MTBC reads were then used for variant calling against the H37Rv reference using Clockwork (v0.12.4) for Illumina, and Clair3 (v1.1.2)/BCFTools (v1.22) for ONT^26–29^. We required two reads to call a variant with Illumina, whilst for ONT we required five. Gnomonicus (v3.1.1) was used to predict resistance according to the second edition of the WHO catalogue (WHOv2) ^18,30^. Uncatalogued mutations in resistance-associated genes were classified as ‘Unknown’, while a result of ‘Fail’ was returned when there were insufficient reads at a genetic loci known to be associated with resistance.

Whether two samples were related was determined by measuring the SNP distance between them, having applied a mask ^17^. The mask excludes repetitive and low complexity regions, as well as regions with frequent errors when mapping Illumina reads ^31^. SNPs within 12 base-pairs of another SNP were also excluded ^32^.

The Clopper-Pearson exact binomial method was used to calculate 95% confidence intervals.

#### ONT Time Subsampling

ONT sequencing generates data continuously, enabling sequencing output to be evaluated at defined time points without interrupting the run. To enable such time-resolved analysis, Min-KNOW was configured to generate per-barcode POD5 files at hourly intervals during sequencing. Following completion of each run, all POD5 files were basecalled and collated to generate FASTQ datasets corresponding to 1, 6, and 72 hours of sequencing. Each time-point dataset was processed independently using the same downstream bioinformatic pipeline as the full 72 hour dataset.

## RESULTS

DNA was extracted from 264 clinical MGIT cultures (Table 2) yielding a median concentration of 12.4 ng*/*µL. The ONT Rapid Barcoding Kit recommends a total DNA input of 200 ng in 10 µL (equivalent to approximately 20 ng*/*µL) for PCR-free library preparation. Based on this criterion, 66% (175/264) of samples fell below the recommended input threshold and were therefore unsuitable for PCR-free library preparation.

**Table 2:** Specimen types (N=264). *Other includes several types such as pleural fluid, bone biopsy, soft tissue and cerebrospinal fluid.

| Specimen Type | Proportion (%) |
| --- | --- |
| Sputum | 78 |
| Bronchoalveolar lavage (BAL) | 10 |
| Pus | 2 |
| Other* | 10 |

Of the 264 samples, 250 met the 1 ng*/*µL internal study cutoff. The remaining 14 samples were excluded from downstream comparative analyses because there was insufficient DNA to perform sequencing on both ONT and Illumina platforms from the same extract.

All 250 samples were successfully sequenced using Illumina, however, five samples failed PCR amplification and were therefore not sequenced using ONT. Among the 245 samples that were successfully amplified for ONT sequencing, the median DNA concentration increased from 13 ng*/*µL to 36.6 ng*/*µL following PCR amplification.

A further five samples were excluded from downstream analyses (see Figure 1) after being identified as non-mycobacterial by both Illumina and ONT sequencing, including *Homo sapiens, Rhodococcus, Pseudomonas aeruginosa, Paenibacillus*, and *Gordonia polyisoprenivorans*.

**Figure 1:**
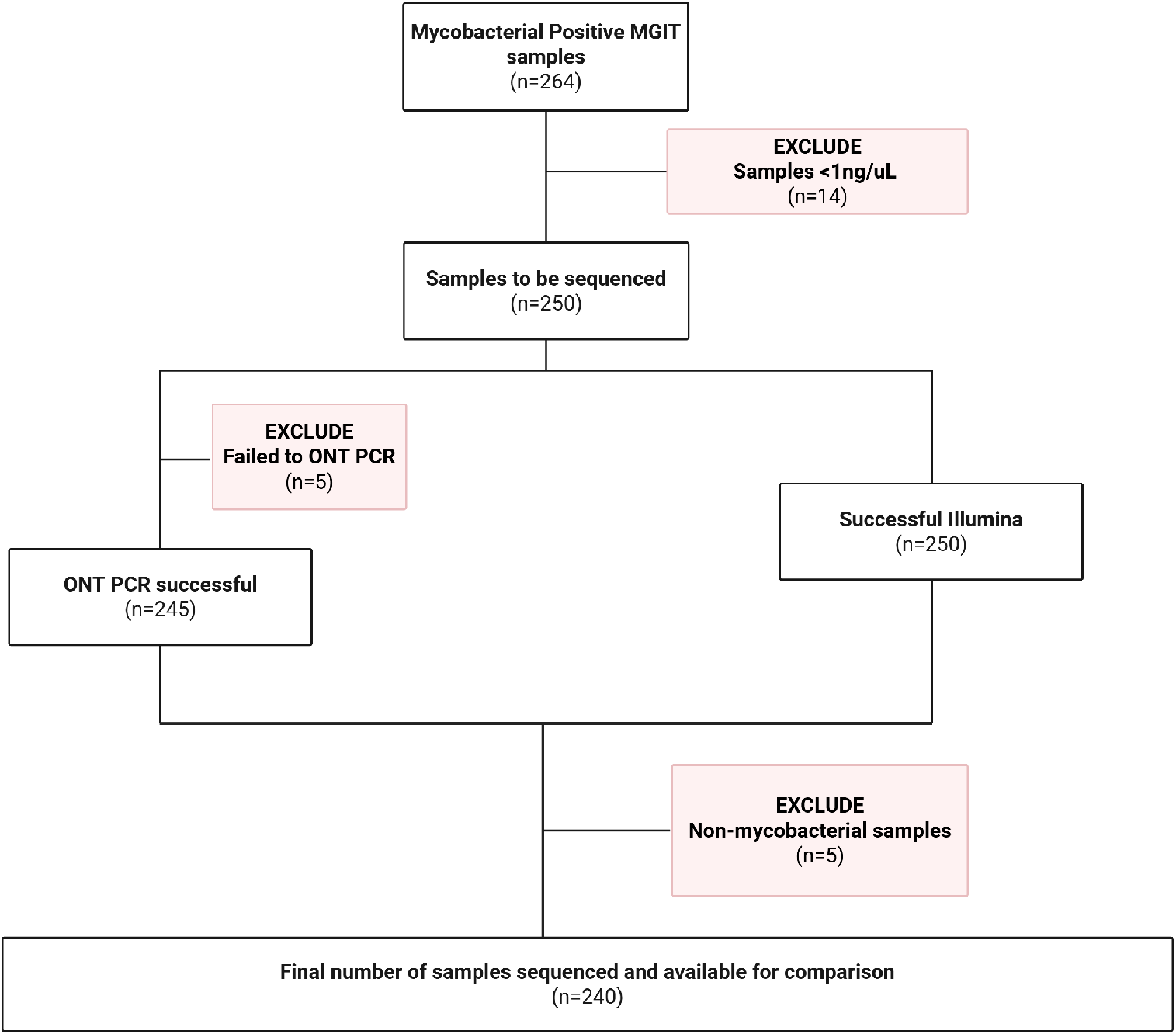
Of the 264 initial samples taken from positive MGIT tubes, 240 were successfully sequenced using both Illumina and ONT platforms and included in downstream analyses. A small number (14) of samples were excluded due to having insufficient DNA, five failed to amplify prior to ONT sequencing and a further five were deemed to not be mycobacterial.

The median sequence yield per sample was 1,342 Mbp for Illumina and 637 Mbp for ONT. For ONT, the median N50 was 3,865 bp, and the median mean read quality score was 16.6 (Figs S1-S4).

### Comparison of species between ONT and Illumina

Of the 240 samples successfully sequenced using both Illumina and ONT, all had at least one *Mycobacterium* species identified by both platforms. Overall, 236/240 (98.3%, 95% CI: 95.8–99.5%) were concordant at the species level. Across the dataset, 21 distinct *Mycobacterium* species were identified.

In three samples, Sylph was unable to assign a species. However, competitive mapping against mycobacterial reference genomes (see Methods) revealed that they had 50-70% genome coverage of *M. gordonae*/*M. paragordonae* in both platforms.

Mixed-species infections were identified in 30 samples by at least one platform. In 26 of these, the same combination of species was consistently detected by both platforms. For these concordant mixed samples, the ratio of Illumina to ONT relative abundance had a median of 0.99 (IQR: 0.93-1.36, Fig. S7).

In the remaining four samples, an additional mycobacterial species was detected by only one platform (Illumina, n=3; ONT, n=1). In two cases, the discordant species was present at low abundance (¡2%), whereas in the other two, it exceeded 30% abundance: *M. abscessus* was detected only by Illumina in one sample, and *M. tuberculosis* only by ONT in another.

Overall, these results demonstrate a high level of concordance between ONT and Illumina for species identification, including in mixed-species samples.

### Platform comparison of *M. tuberculosis* samples

A total of 46/240 (19.2%) samples were determined to contain *M. tuberculosis* by both platforms. Four samples were excluded due to low read depth when sequenced by either Illumina (n=3) or both platforms (n=1), leaving 42 *M. tuberculosis* samples each with a pair of high-quality genomes included in downstream analyses.

The same sub-lineage was assigned across platforms for all 42 samples. The samples were phylogenetically diverse with all four major lineages represented and a total of 20 distinct sublineages (Fig. S8) identified.

Very few mutations associated with antimicrobial resistance were detected in this dataset. With the exception of delamanid, only four samples showed resistance to the 15 drugs covered by WHOv2 (Fig. 2). In three samples, the detected mutation was identical in both platforms, but one sample had two mutations triggering resistance to the same drug: one identical across platforms and one unique to Illumina (a minor SNP with 15% read support). Around half (23/42, 54.8%) of the ONT genomes were classified as resistant to delamanid due to a 62 bp deletion that starts seven bases before the end of *fbiC*. According to WHOv2, this deletion triggers a loss-of-function rule resulting in a ‘Resistant’ call. However, this region corresponds to a tandem repeat downstream of *fbiC*, and the deletion is not expected to affect the primary coding sequence (Supplementary §5.2).

**Figure 2:**
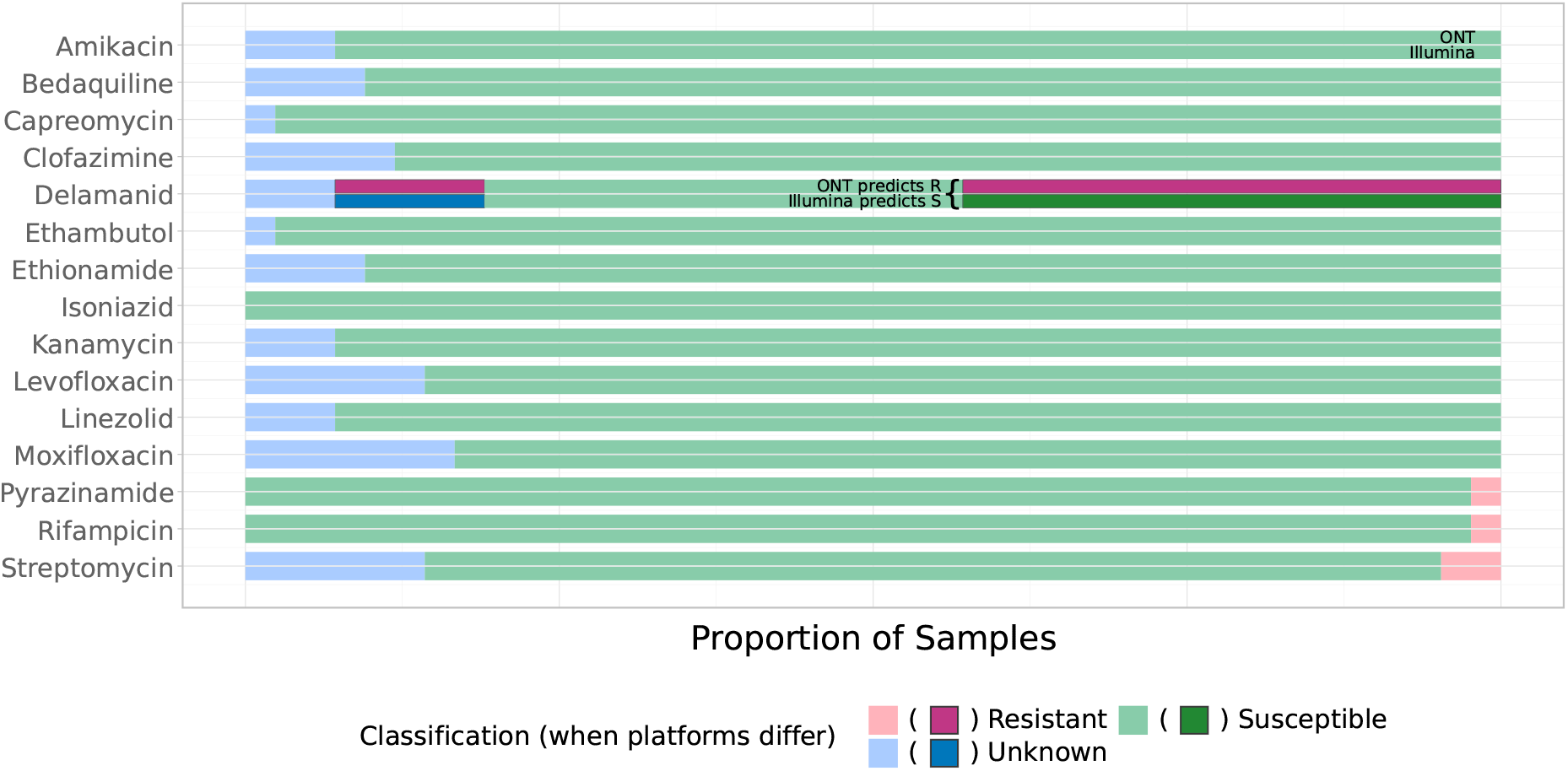
Comparison of genotypic resistance classification between ONT and Illumina. For each drug the lower bar shows the proportions of samples classified as Unknown/Susceptible/Resistant by Illumina, and the top bar by ONT for the corresponding samples. Where platforms differ the sections are shown in bold with darker colours. i.e. The annotated part of Delamanid shows that 43% of samples were considered susceptible by Illumina but resistant by ONT. All drugs except Delamanid have complete agreement between platforms.

All samples had sufficient depth at all genetic loci associated with resistance (Group 1) according to WHOv2, and no samples returned a ‘Fail’ result for any drugs. The pipeline returned a result of ‘Unknown’ for uncatalogued mutations in resistance genes. These were entirely concordant between ONT and Illumina (38 mutations across 23 samples). We also found 417 mutations not associated with resistance. Most (409/417, 98.1%) were common to both platforms with the remaining mutations being SNPs belonging to a minority population; five unique to Illumina, two unique to ONT, and one indel unique to ONT.

Genomic similarity was assessed using SNP distance (see Supplementary §5.3 for details of calculation and masking). We first assessed SNP distances between ONT and Illumina genome pairs for each of the 42 *M. tuberculosis* samples (Fig. 3). The mean distance was 0.3 SNPs with 34/42 (81.0%) samples having a zero distance and therefore being identical. The largest distance was for a single sample with seven SNPs between ONT and Illumina.

**Figure 3:**
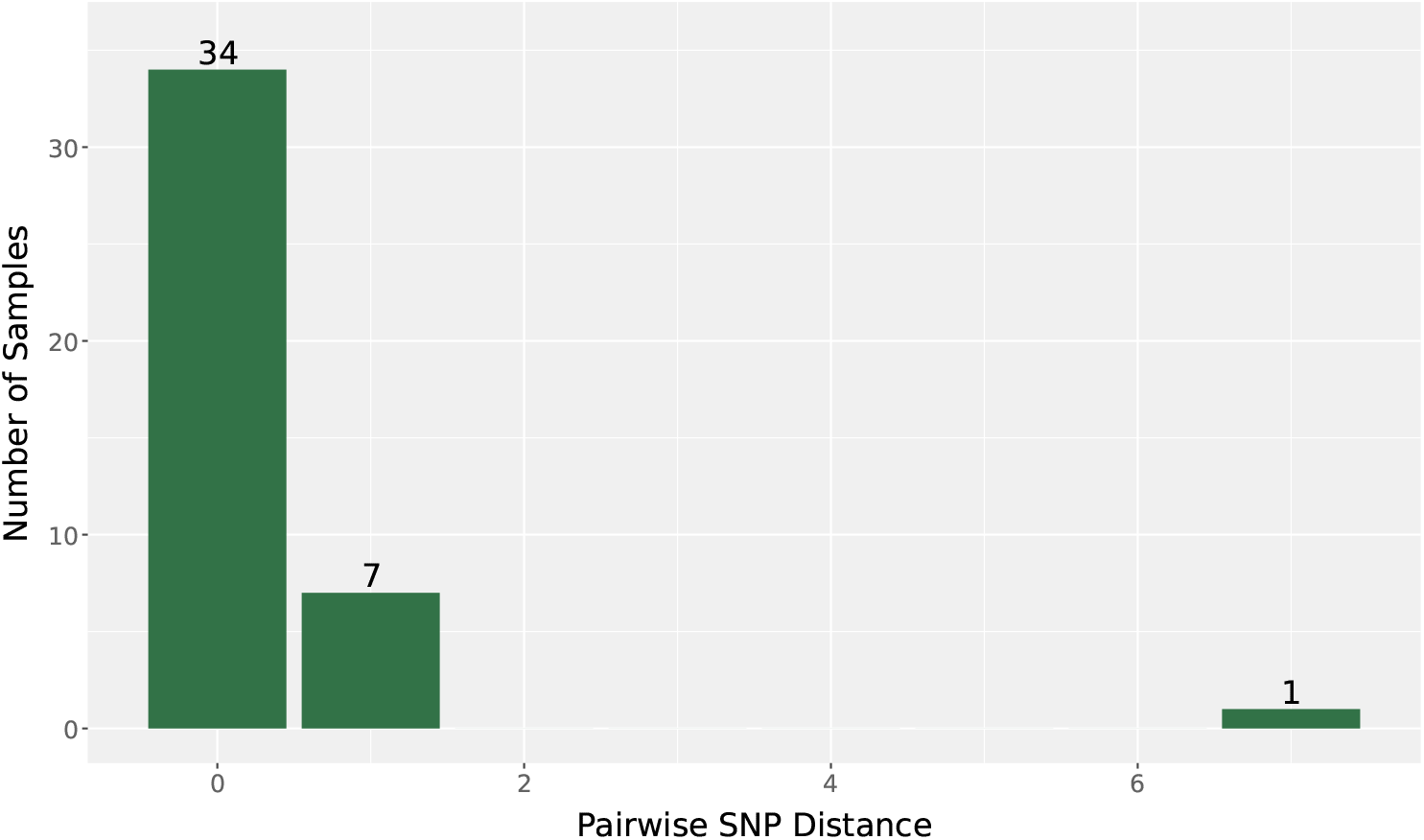
Bar chart of the measured SNP distance when the same sample is sequenced by both Illumina and ONT platforms. Only the 42 *M. tuberculosis* samples are considered, hence 34/42 (81.0%) of samples are identical from the perspective of detected SNPs.

Three pairs of samples originated from the same patients at different time points. Within each pair, all four genomes (two samples, each having been sequenced using Illumina and ONT) were expected to be highly similar. All combinations of genome pairs were identical (zero SNPs) regardless of which mix of platforms was used to sequence them.

For all pairs of samples we calculated the absolute difference in SNP distance between platforms as a fraction of the Illumina-distance (excluding the three zero-distance pairs described above). The median of this metric across all pairs was 0.7%, or 0.8% if considering cross-platform comparisons (Supplementary §5.3).

Overall, ONT and Illumina demonstrated highly concordant performance for lineage assignment, SNP-based relatedness, and resistance detection

### ONT sequencing after 1, 6 and 72 hours

To assess the reliability of early ONT sequencing data, we analysed the reads generated after 1, 6, and 72 hours. Subsampling of reads at 1 hour and 6 hours demonstrated high concordance with species assignments obtained from the final 72 hours dataset. Of the 240 samples, 237 had a species identified after 72 hours. Of these, 224/237 (94.5%) were correctly identified after 1 hour, increasing to 233/237 (98.3%) after 6 hours (Fig. 4, S10). The remaining four samples at 6 hours differed only by the absence of low-abundance secondary species (¡1%) that were detected after 72 hours.

**Figure 4:**
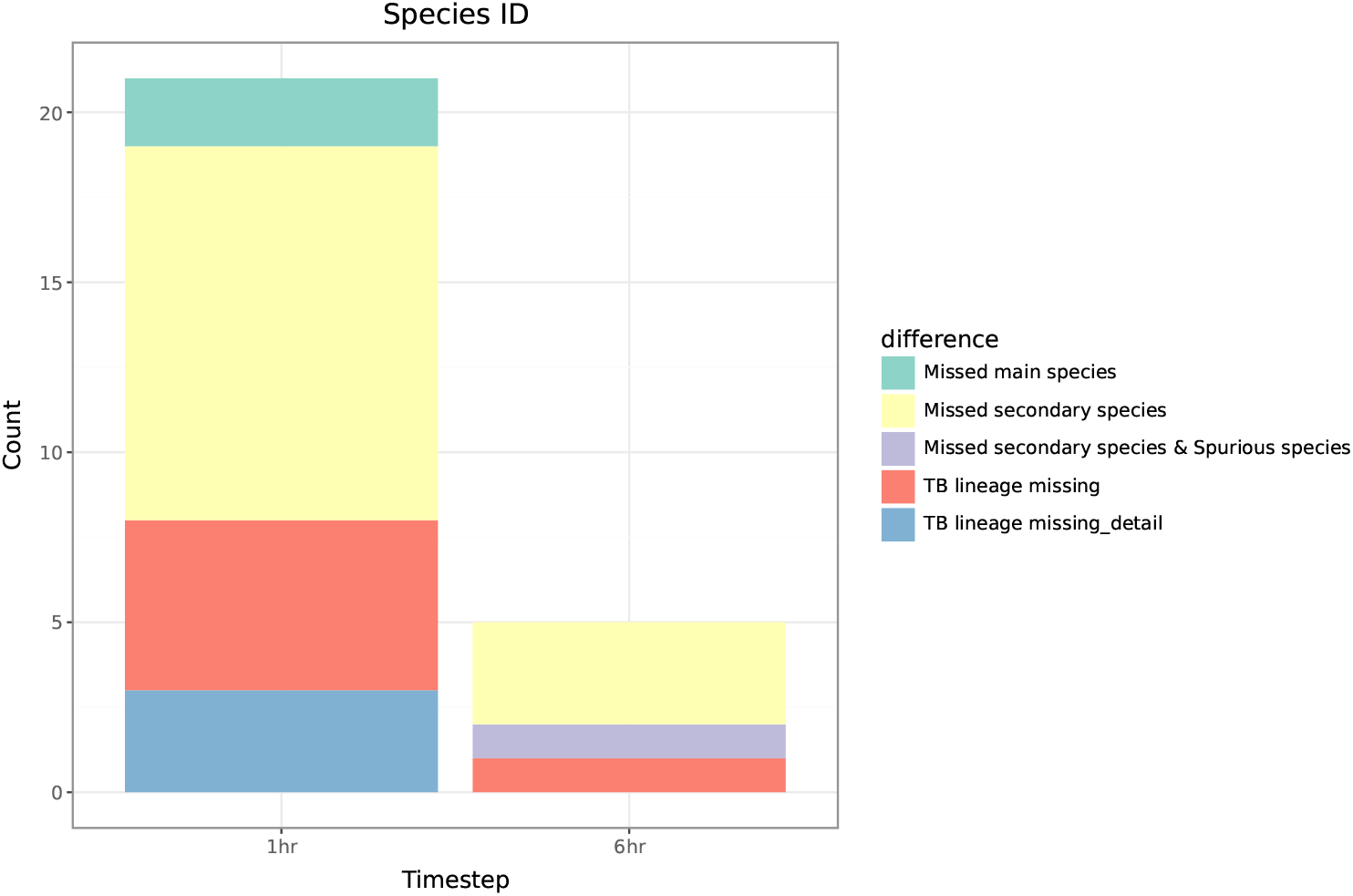
The differences in the species identified and lineage classified after 1 and 6 hours of sequencing compared to the result after 72 hours. We are therefore assuming that the result after 72 hours is correct. At 1 hour 216/237 (91.1%) samples were concordant with 72 hours. This increased to 232/237 (97.9%) at 6 hours.

In two samples, no species was identified after 1 hour due to insufficient reads, causing early pipeline termination. There were no instances of incorrect primary species assignments at 6 hours, and only one secondary species was detected that was not present in the 72 hour dataset; which corresponded to a non-mycobacterial organism.

Lineage assignment for *M. tuberculosis* was performed using Mykrobe. After 1 hour, eight samples showed discordance compared to the 72 hour result, including missing lineage calls or reduced sub-lineage resolution. By 6 hours, this was reduced to a single discordant sample (Fig. 4).

Sequencing depth increased substantially over time. After 1 hour, mean read depth for the dominant species ranged from 0.1–10× (Fig. S11), with genome coverage varying between 20–100% (Fig. 5). After 6 hours, mean read depth increased by approximately an order of magnitude, resulting in improved genome coverage. Further sequencing to 72 hours increased read depth but provided minimal additional gains in genome coverage for the dominant species.

**Figure 5:**
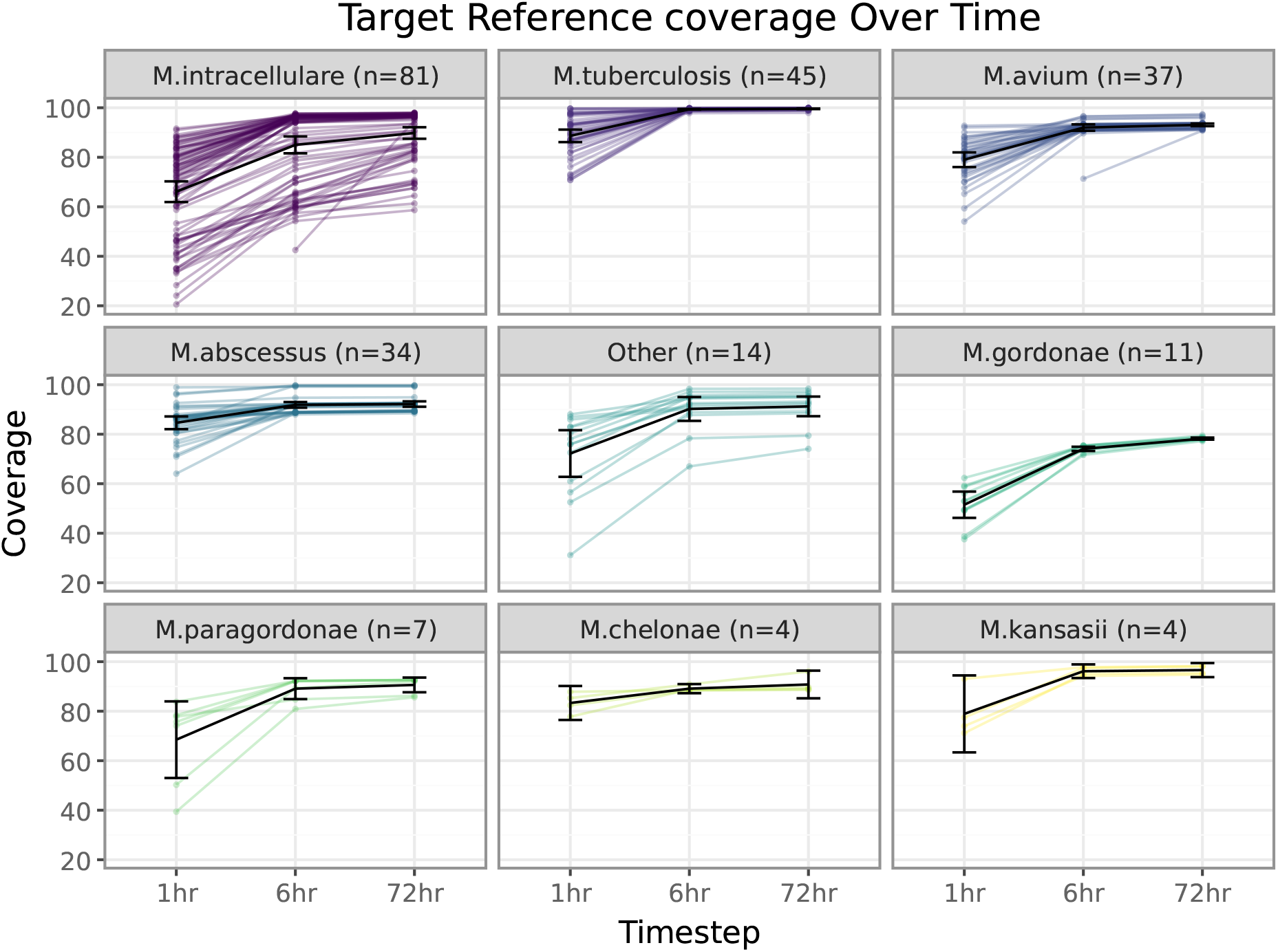
Genome coverage of major species after 1, 6, and 72 h of sequencing. *M. gordonae* had markedly lower coverage even after 72 h, suggesting that the reference poorly described our samples.

For *M. tuberculosis*, mean read depth increased from 21.1 *±* 12.9 at 6 hours to 142.9 *±* 86.1 at 72 hours. However, substantial variability between samples indicates that while some can be fully analysed after 6 hours, others require longer sequencing to achieve sufficient depth for downstream analyses.

## DISCUSSION

This study demonstrates that, with optimized DNA extraction from MGIT cultures and a modified PCR-based ONT library preparation protocol, long-read sequencing of mycobacteria can achieve results comparable to those obtained using Illumina sequencing. The workflow generated sufficient DNA input to enable reliable ONT sequencing from primary MGIT cultures, resulting in low failure rates and accurate species identification. For *Mycobacterium tuberculosis* samples, lineage assignment, antimicrobial resistance prediction, and identification of related isolates were also highly concordant between platforms, supporting the feasibility of PCR-amplified ONT sequencing for routine genomic characterization of MGIT-derived isolates.

The semi-automated extraction workflow reduced manual handling steps, a recognized bottleneck in mycobacterial sequencing pipelines. The Rapid PCR Barcoding approach overcame the frequent limitation of low DNA yield from MGIT cultures and is not only advantageous in this setting but also broadly applicable to other low-concentration or low-input DNA samples, helping to address a long-standing barrier to long-read sequencing. The high proportion of samples falling below the recommended DNA mass input threshold for the Rapid Barcoding Kit further supports the use of PCR-based amplification for MGIT-derived DNA, where yield and purity are frequently limiting. In this study, a pragmatic internal threshold of 1 ng/µL was used to ensure sufficient material for both ONT and Illumina sequencing; however, ONT Rapid PCR Barcoding can be performed with substantially lower DNA inputs (as low as 1 ng in 3 µL, equivalent to around 0.33 ng/µL, according to the manufacturer’s instructions), indicating that samples below our threshold could still be successfully sequenced using a single-platform approach. Together, these optimisations address a key practical barrier to the adoption of ONT sequencing in routine mycobacterial workflows.

Despite recent progress, ONT reads still have higher error rates than Illumina reads. We observed, however, that the discrepant SNPs in ONT data clustered within discrete genomic regions rather than being randomly distributed. This pattern suggests that selective masking or region-aware filtering could improve cross-platform concordance, particularly for phylogenetic applications. Importantly, the magnitude of disagreement outside these regions was small, indicating that ONT-derived genomes are sufficiently accurate for species identification and lineage assignment. Continued bioinformatic refinements, particularly to variant calling, are likely to further enhance cross-platform reproducibility, enabling reliable comparison of results between laboratories using different sequencing technologies, consistent with findings from recent comparative studies ^17^.

Two samples showed evidence of mixed mycobacterial species. In one case, *M. tuberculosis* was detected by ONT but not by Illumina, whereas in the other *M. abscessus* was detected by Illumina but not by ONT. Given the magnitude of these discrepancies and the otherwise high concordance observed between platforms, these findings are most plausibly explained by sample-level contamination or biological heterogeneity rather than systematic sequencing or bioinformatic differences. Such events are well recognised in routine mycobacterial workflows and highlight the importance of careful sample handling, result interpretation, and repeat testing, particularly when low-abundance species are present.

A subset of Illumina datasets contained *>*5% non-mycobacterial reads by Sylph taxonomic profiling after Kraken2 filtering. This likely reflects differences in how short and long reads interact with taxonomic classifiers. Longer ONT reads provide greater genomic context and therefore are more confidently assigned by tools such as Kraken2, whereas shorter Illumina reads are more frequently left unclassified rather than misclassified. These unclassified reads are subsequently assigned low-confidence labels by downstream tools such as Sylph in our pipeline, inflating the apparent proportion of non-mycobacterial sequences. This highlights the need for harmonized, platform-aware taxonomic approaches when comparing sequencing modalities.

In a clinical laboratory context, ONT offers several practical advantages. Data is generated continuously, enabling real-time monitoring of sequencing runs and allowing runs to be terminated once sufficient coverage has been achieved. Subsampling analyses demonstrated that accurate species identification and lineage assignment were frequently achievable after six hours of ONT sequencing, although variability in sequencing depth meant that longer runs were required for antimicrobial resistance prediction in some samples. This variability underscores the need for sample- and application-specific sequencing quality thresholds in diagnostic settings. Perhaps more importantly, this protocol has the potential to bring downstream genomic characterisation in-house into routine clinical diagnostics, reducing reliance on outsourcing to reference laboratories and enabling more timely access to genomic results.

This study has limitations. It was performed on MGIT-positive cultures rather than primary clinical specimens and therefore cannot replace culture or direct diagnostic testing; instead, it complements them by accelerating downstream genomic characterization. Illumina sequencing was performed by a commercial provider whereas ONT sequencing was conducted in-house, which may introduce methodological differences. Resistance prediction for NTM was not evaluated, and antimicrobial resistance diversity in the *M. tuberculosis* samples was limited, although there was good coverage of resistance conferring genes. A small proportion of samples (5/250; 2%) failed PCR amplification, highlighting the potential need for repeat extraction or alternative amplification strategies for challenging specimens. In addition, variability between ONT sequencing runs was observed, with one flow cell producing reduced throughput and read quality, underscoring the importance of run-level quality control thresholds. Further evaluation in larger and more diverse cohorts will be required prior to routine diagnostic implementation.

In conclusion, by addressing the challenges of low-yield MGIT extractions and establishing a reliable PCR-based ONT workflow, this study demonstrates the feasibility of long-read sequencing as an alternative to established short-read technologies for mycobacterial genomics, supporting its further validation in routine laboratory settings.

## ACKNOWLEDGMENTS

We thank EIT Oxford for deploying our pipeline in their cloud platform and to ORACLE Corporation for access to their cloud.

## AUTHOR CONTRIBUTIONS

CSB - Conceptualization, Data curation, Investigation, Methodology, Project administration, Resources, Validation, Writing - original draft, Writing - review and editing. MC - Data curation, Formal analysis, Methodology, Validation, Visualisation, Writing - original draft, Writing - review and editing. JG - Methodology, Investigation, Resources, Validation. AH - Methodology. ER - Investigation. HW - Methodology. BD - Investigation, Resources. RC - Investigation, Resources. HT - Resources, Software. JW - Software. RT - Software, Writing - review and editing. TEAP - Conceptualization, Writing - review and editing. PWF - Software, Writing - review and editing. MM - Methodology, Investigation, Resources, Validation. DWC - Conceptualization, Supervision, Writing - review and editing.

## DATA AVAILABILITY STATEMENT

Reads will be deposited in the ENA (PRJEB106303) on acceptance and the project code added at proof.

## ETHICS APPROVAL

MGIT samples growing Mycobacteria were collected from the NHS microbiology laboratory of Oxford University Hospitals (OUH) NHS Foundation Trust and residual culture medium were collected. This was approved by the Oxford Research Ethics Committee REC Reference 17/LO/1420 IRAS project ID 228657.

