## supplementary_material for "Validation of an optimised Oxford Nanopore sequencing workflow versus Illumina for mycobacteria from primary MGIT culture"

### 1 Description of supplementary materials

The following supplementary tables have been provided:

1. The mycobacterial references used in the competitive mapping step
2. Species classification (Sylph) and lineage assignment (Mykrobe) per sample
3. AMR predictions
4. All mutations found in the samples
5. SNP distance between ONT and Illumina assembly for the same sample
6. All pairwise distances with all four permutations of technologies
7. Species frequency counts according to Sylph for both platforms
8. Results from competitively mapping reads against reference genomes.

### 2 Read statistics

All statistics here have been calculated post human read removal. Points are coloured by proportion of reads that were determined as human, which was the cause of low yield samples in both Illumina and ONT. Additionally, flow cell 7 on ONT performed substantially worse than the others in terms of overall yield.

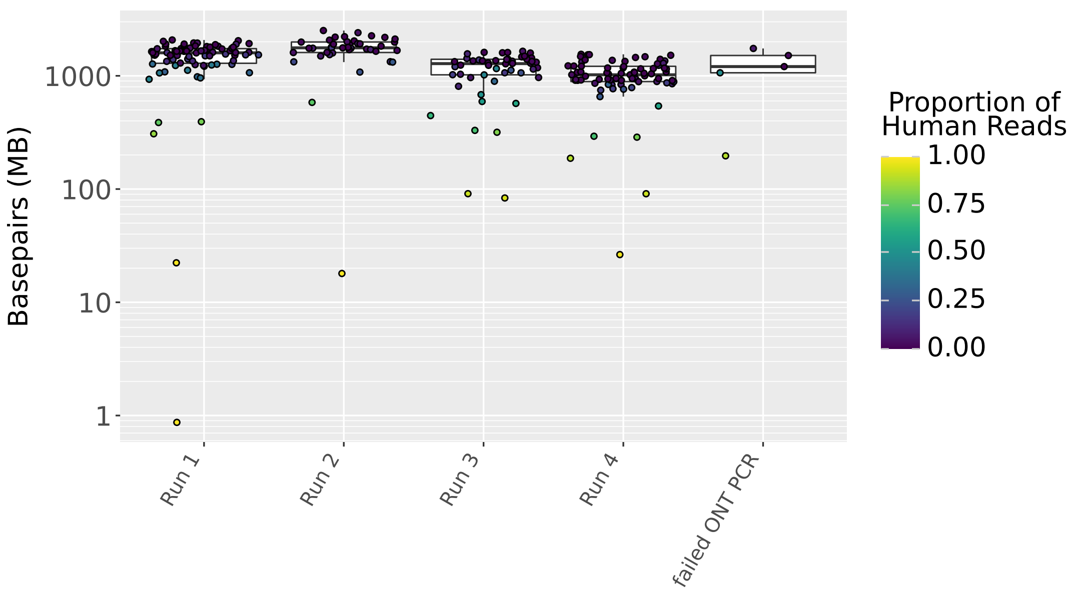


Figure S1: Illumina sequencing yield per sample. The five samples which failed to PCR for ONT are shown separately.

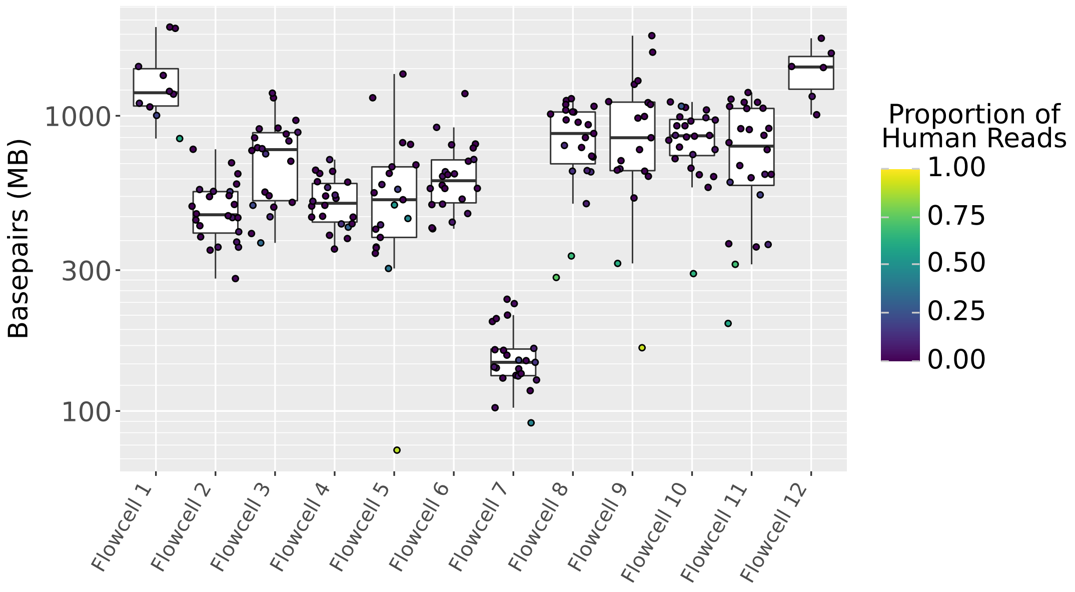


Figure S2: ONT sequencing yield per sample

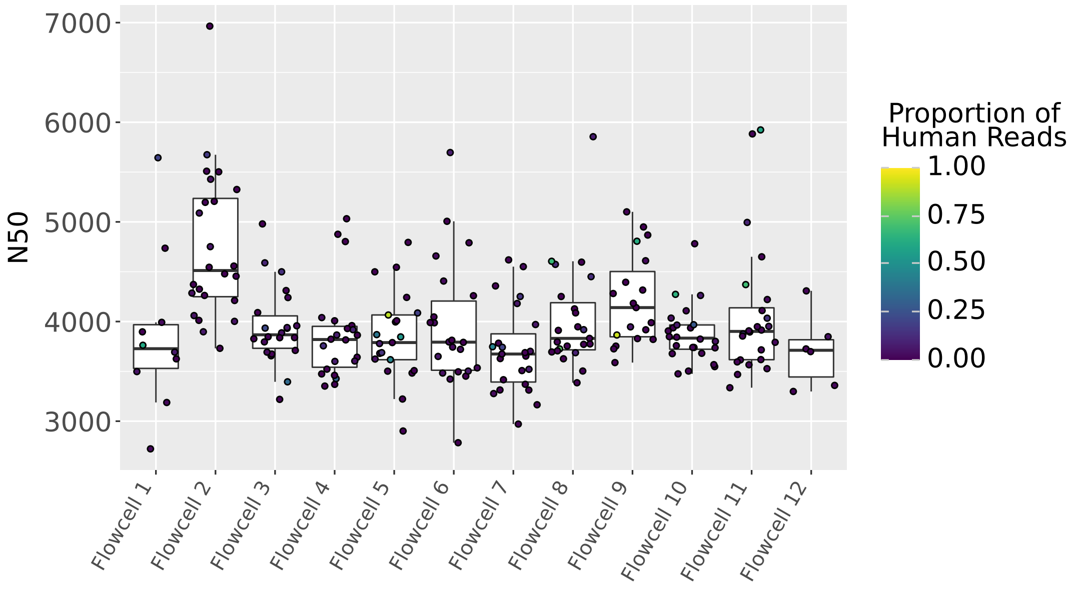


Figure S3: ONT read length N50 per sample

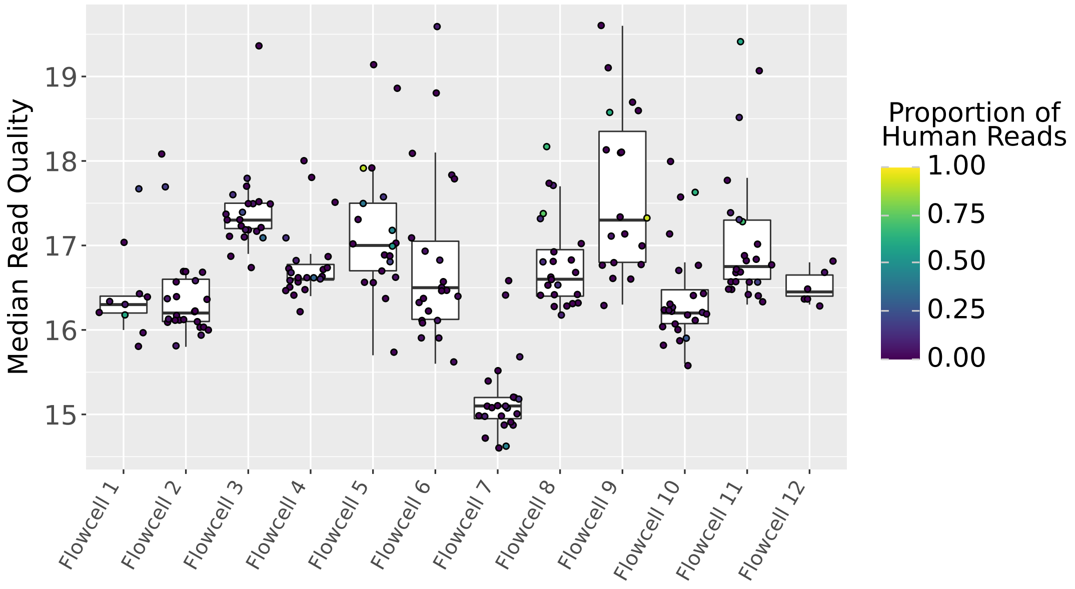


Figure S4: ONT median read quality per sample

#### 3 Sylph thresholds for species comparison

Sylph is a k-mer based tool for species identification from sequencing reads. It is designed to be sensitive to potentially very small quantities of a species. This produced some complications with comparing results at very small proportions (<1% taxonomic abundance).

Within our three control samples for ONT we still found reports for minor species. The worst case being a *M. avium* call in the BCG control which had 0.14% taxonomic abundance and estimated depth of 0.32. To avoid issues with spurious minor calls we required at least 0.5% taxonomic abundance and 0.5 depth to report a difference in species ID. Figures S5, S6 shows the distribution of species calls identified at 72 hours but not 1 or 6 hours. These minor calls may be a result of genuine contamination, barcode crossover or bioinformatic error.

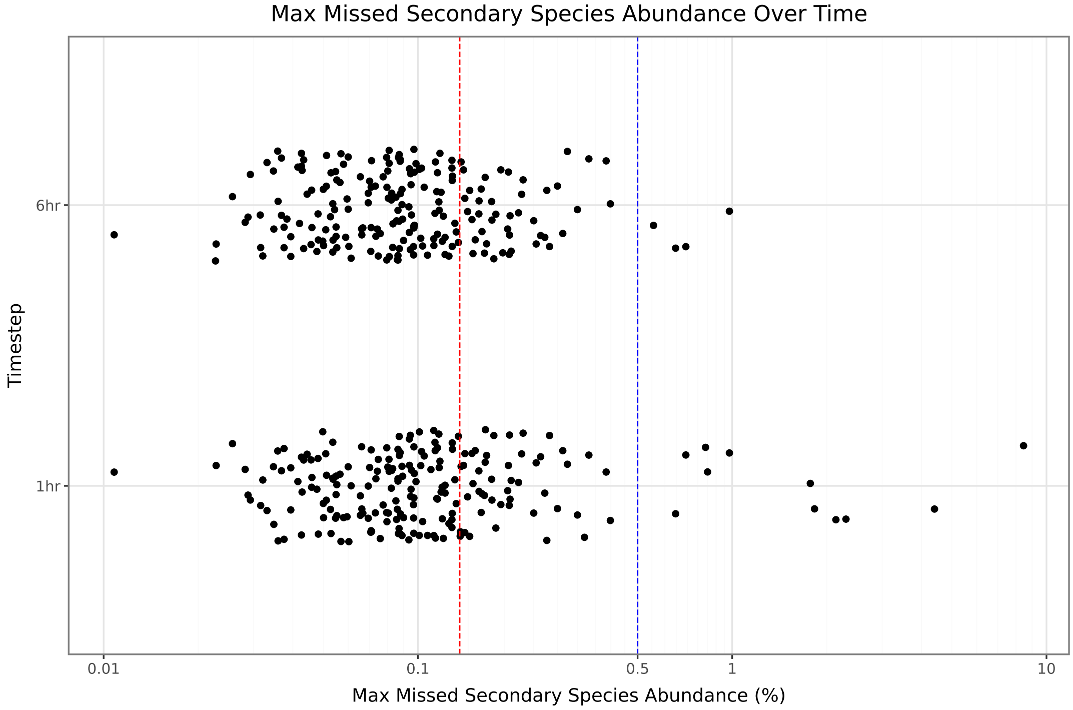


Figure S5: Taxonomic abundance of missed species calls. For each sample the highest abundance species call at 72 hrs not identified at 1 or 6 hrs is plotted. The red line marks the BCG control and the blue line the 0.5 filter used.

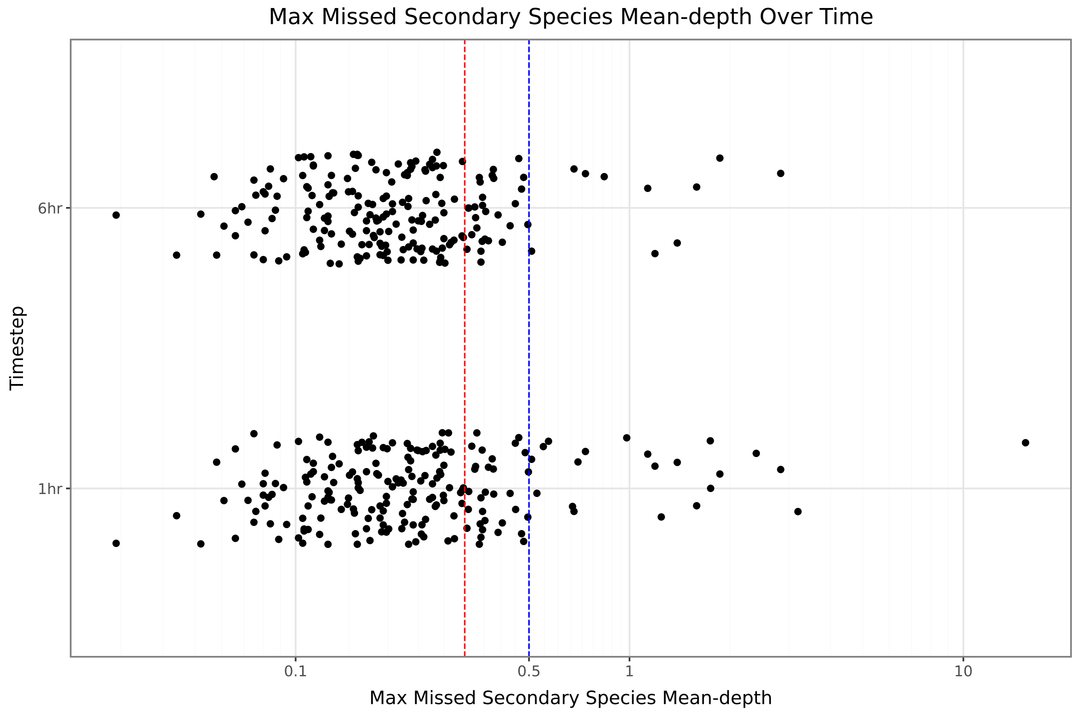


Figure S6: Estimated depth of missed species calls. For each sample the highest depth species call at 72 hrs not identified at 1 or 6 hrs is plotted. The red line marks the BCG control and the blue line the 0.5 filter used.

### 4 Detection of mixed infections

In 30 samples more than one mycobacterial species was detected (using a threshold of 0.5% taxonomic abundance from Sylph) by at least one of the platforms. In two samples there were three species detected, whilst the remaining 28 there were two. For these samples we considered the primary species to be the one with the highest minimum of Sylph taxonomic abundance between the two platforms, and called the rest secondary.

For each of these secondary species we compared the abundance estimate between Illumina and ONT. Species were marked as concordant if they were identified in both platforms even if abundance was below 0.5% in one of the platforms (Fig S7).

For the concordant secondary species the ratio of ONT abundance to Illumina abundance had a median of 0.99 (IQR: 0.93-1.36).

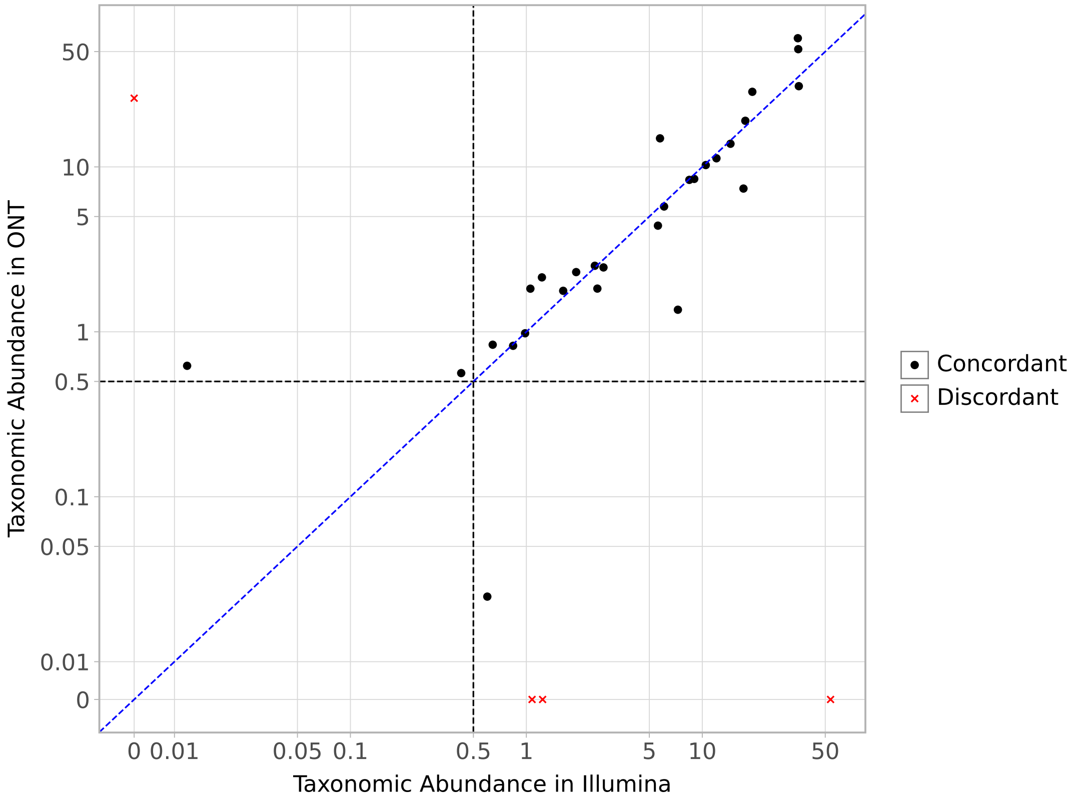


Figure S7: Taxonomic abundance estimates for secondary mycobacterial species. The blue diagonal line shows $y = x$ and the black lines indicates the minimum abundance required for a species to be counted.

### 5 Comparison of *M. tuberculosis* samples

#### 5.1 Lineage assignment

Of the 46 samples identified as TB, three had low depth in Illumina and one had low depth in both platforms. Mykrobe was used to assign lineages and the only sample with a discrepancy had 1.1.2 in ONT and 1.1 in Illumina (which had low depth). Note that Mykrobe lineage calls with the filter `LOW_PERCENT_COVERAGE` were excluded as having low support as they were only seen in very low depth samples. Lineage 4 was the most common in our collection (18 samples), followed by Lineages 3 (15), 1 (6), 2 (2) and *M. bovis* (1). Within those lineages, 20 distinct sub-lineages were identified.

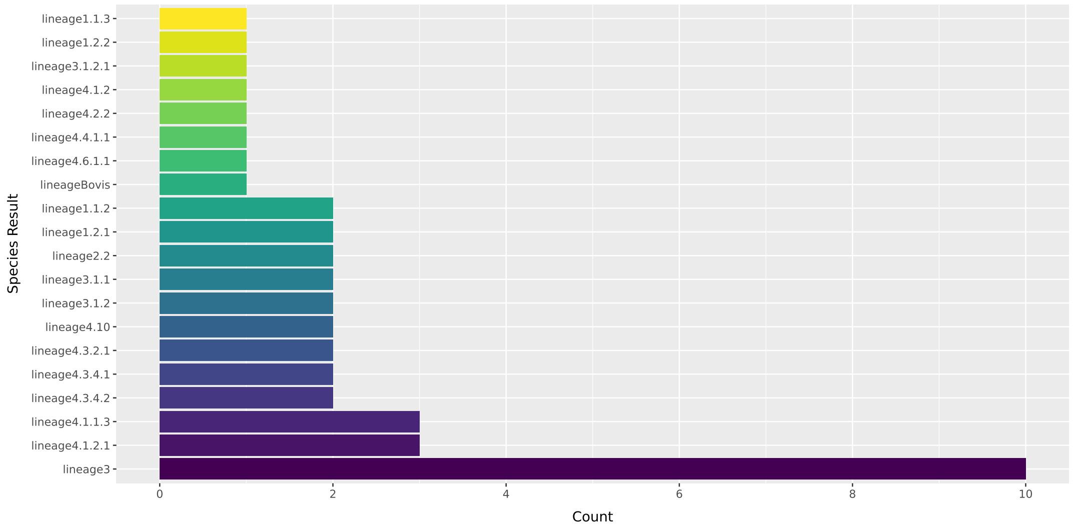


Figure S8: All four main lineages were identified with Lineage 4 being the most common.

#### 5.2 Detection of antimicrobial resistance (AMR)

##### 5.2.1 Deletions in the *fbiC* gene

The end of the *fbiC* gene is a tandem repeat with a 62bp section of DNA which occurs twice in full and finally a section of 43bp. In 23 of our samples, the ONT assemblies had one of these repeats deleted. Since each 62 bp section contains the *fbiC* stop codon, the deletion of just one of the repeats does not therefore alter the resulting primary protein sequence.

#### 5.3 Relatedness

Whether two samples were related was determined using SNP distance. This was calculated by comparing the fixed-length assemblies and counting the number of sites with differing base calls. If a site had a null call in either genome then it was excluded. Further, a mask was used to further exclude a subset of sites known to be repetitive, low complexity or have poor Illumina mapping[1,2]. Fig. S9 shows how certain sites are hotspots for differences between the two platforms, most often with ONT having a SNP, relative to H37Rv, that is not found by Illumina.

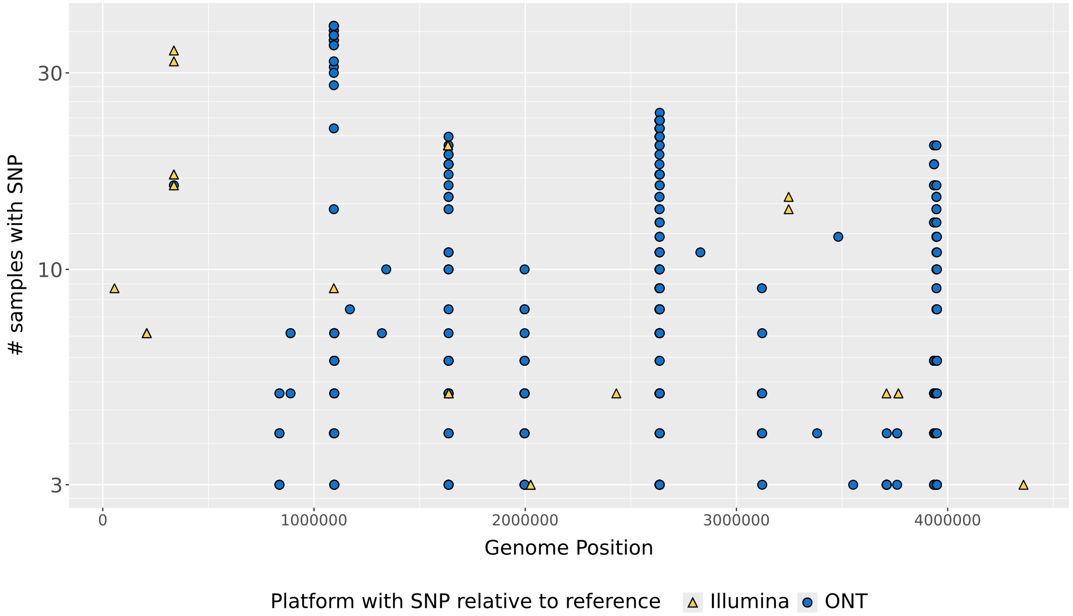


Figure S9: Manhattan plot showing distribution of SNPs across the genome between the Illumina and ONT assemblies. The y-axis indicates the number of samples that had a SNP at that site between the two platforms. The shape/colour indicates which platform had the SNP relative to the H37RV reference.

For evaluating the difference in SNP distance between platforms we define the following metrics of difference in SNP distance ($O$ is ONT, $I$ is Illumina).

$$\text{dist}_{u,v}(a,b) = \text{distance between u and v using platforms a and b respectively.}$$

$$\text{divergence}_{u,v} = \frac{|\text{dist}_{u,v}(O,O) - \text{dist}_{u,v}(I,I)|}{\max(\text{dist}_{u,v}(I,I),1)}$$

$$\text{complex divergence}_{u,v} = \frac{|\max_{a,b \in O,I} \text{dist}_{u,v}(a,b) - \min_{a,b \in O,I} \text{dist}_{u,v}(a,b)|}{\max(\text{dist}_{u,v}(I,I),1)}$$

We can calculate these metrics across all pairs of samples. The mean divergence is 1.1% (median 0.7%) and the complex divergence is 1.4% (median 0.8%).

#### 6 ONT time subsampling

For 3/240 samples, no mycobacterial species could be identified using Sylph after 72 hours (these are the likely *M. gordonae*/*M. paragordonae* samples as assigned by competitive mapping). These have therefore been excluded from this analysis.

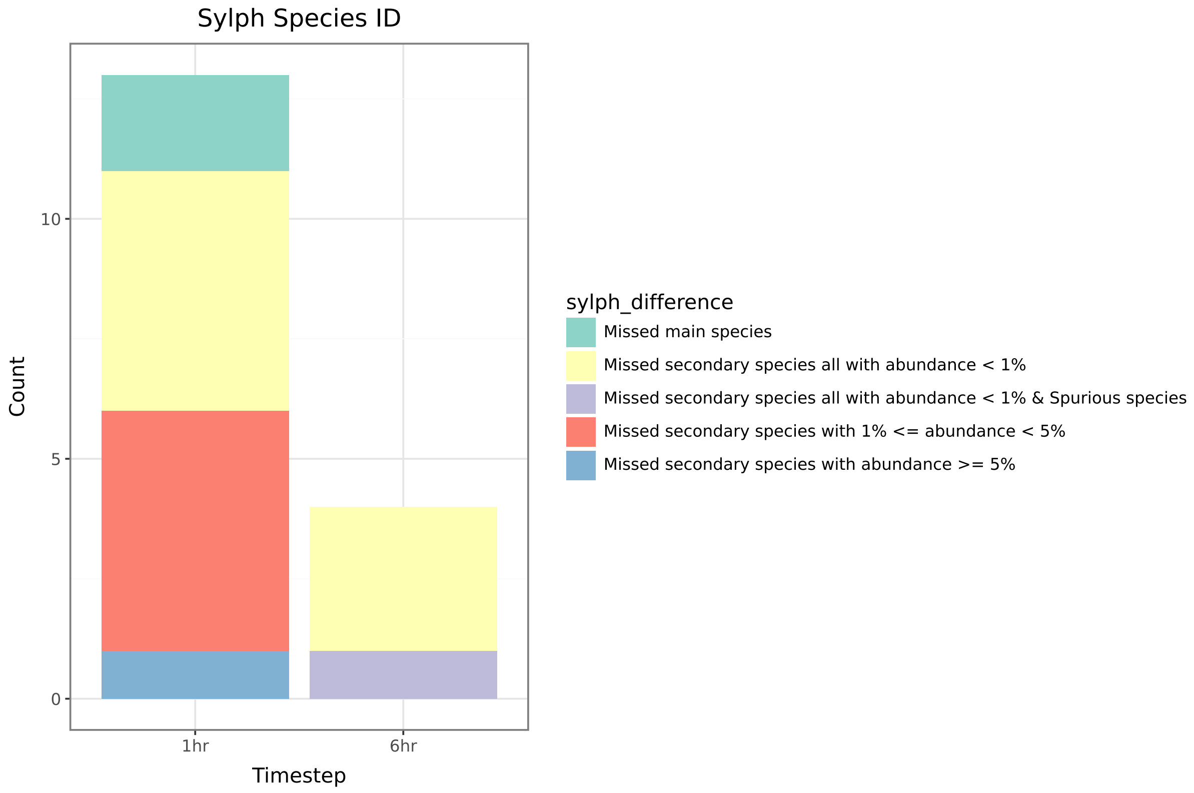


Figure S10: Sylph species identification differences at 1 and 6 hrs compared to 72 hrs for the 237 samples. A "spurious species" is a species not present after 72 hrs sequencing that was identified as being present at one of the earlier two timepoints. The main species was the highest abundance species at 72 hrs. At 1 hr there are 224 concordant samples, increasing to 233 at 6 hrs.

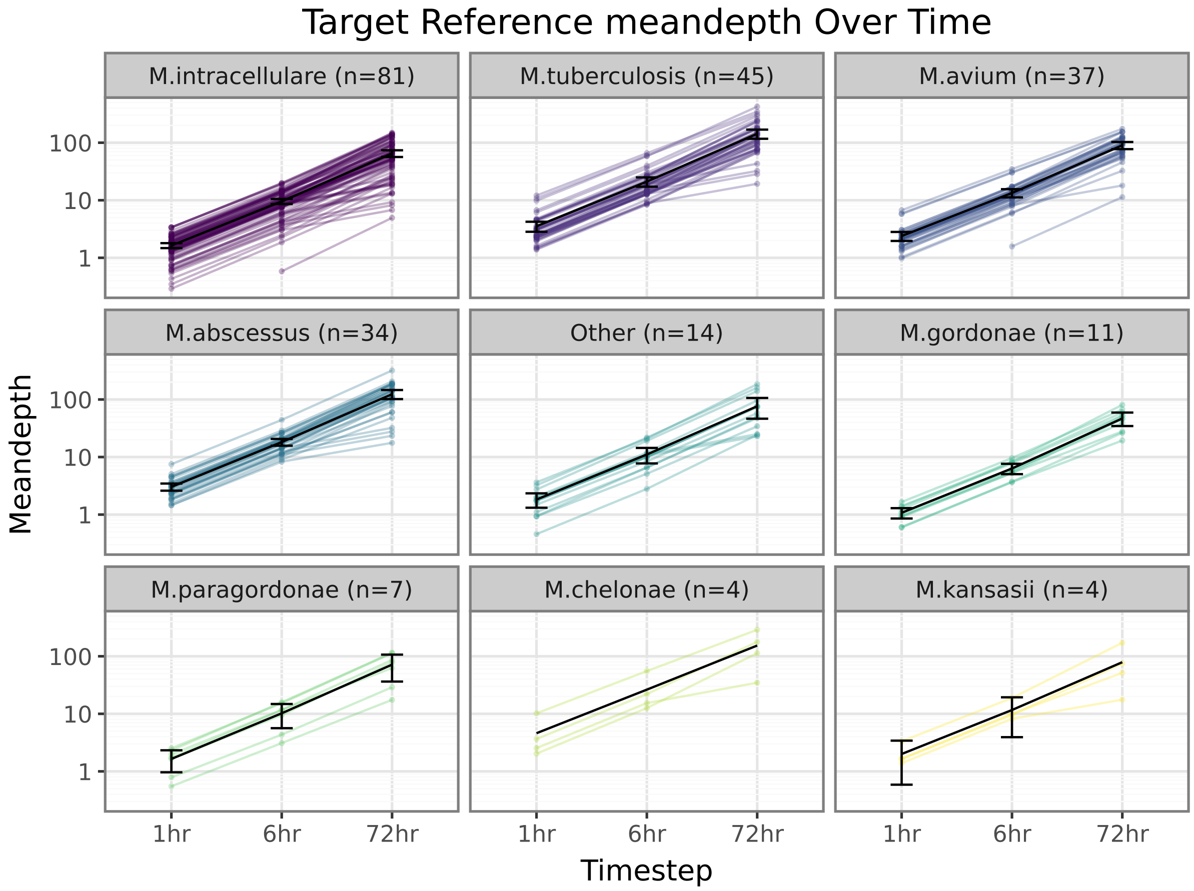


Figure S11: The mean read depth of the major species identified increases with the length of the sequencing run. Error bars show the 95% confidence internal (where non-negative due to the log scale)
